## supplementary files for "TGFβ blocks IFNα/β release and tumor rejection in spontaneous mammary tumors"

a

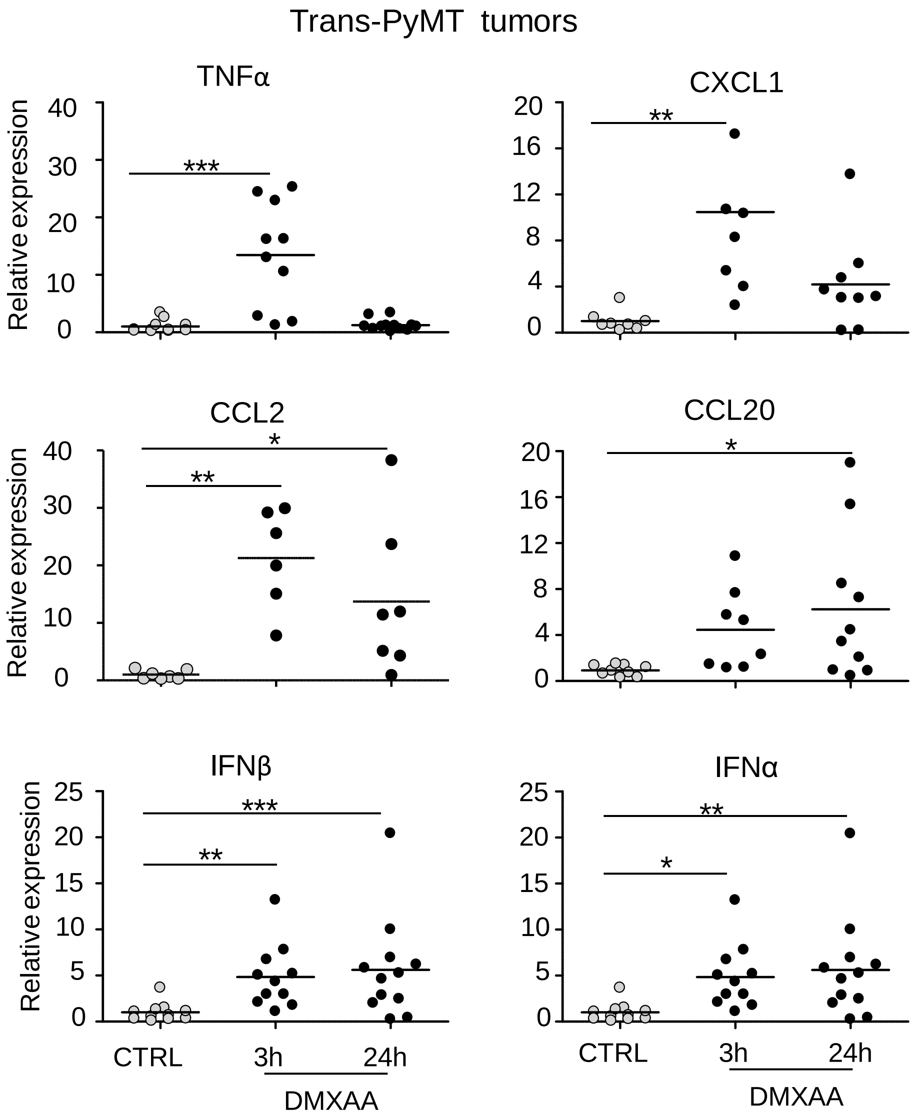
Trans-PyMT tumors
TNFα
CXCL1
CCL2
CCL20
IFNβ
IFNα
Relative expression
CTRL
3h
24h
DMXAA

b

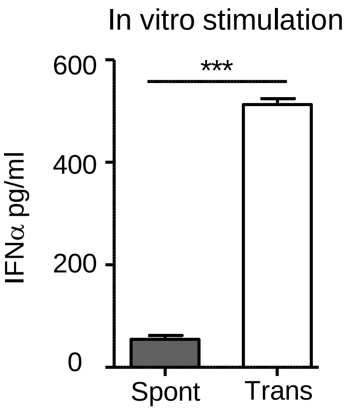
In vitro stimulation
IFNα pg/ml
600
400
200
0
***
Spont
Trans

FigS1

**Figure S1. Expression of type I IFN, TNFα and chemokines in tumors of Trans-PyMT mice induced by DMXAA. (a)** Trans-PyMT mice were injected i.p. with DMXAA or DMSO, and mRNA levels of cytokines and chemokines were measured as indicated in Fig. 2a. Cumulative data of 7-11 mice, from 3-5 independent experiments are shown. Tukey's Multiple Comparison Test. **(b)** Tumors from Spont- and Trans-PyMT mice were dissociated and restimulated *in vitro* with DMXAA (250µg/ml) to measure IFNα production in the supernatant after 24h by ELISA as described in Fig. 5. Results, expressed as mean ± s.e.m. (Spont-PyMT mice: n=3, Trans-PyMT mice: n=7; from 6 independent experiments) were analyzed with a Student *t*-Test.

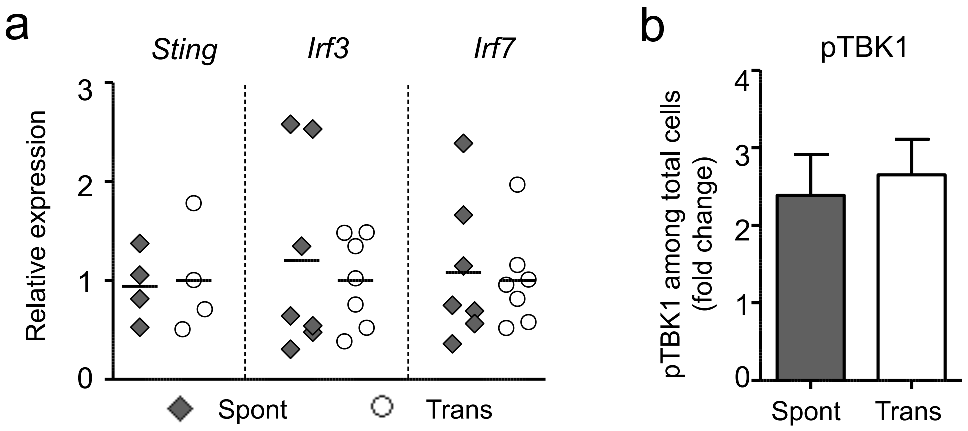
a
Sting
Irf3
Irf7
Relative expression
0
1
2
3
Spont
Trans
b
pTBK1
pTBK1 among total cells (fold change)
0
1
2
3
4
Spont
Trans

FigS2

**Figure S2.** **(a)** Expression of *Sting*, *Irf3* and *Irf7* mRNA transcripts in tumors of Spont-and Trans-PyMT mice**.** The relative expression is shown, normalized to Trans-PyMT tumors. Spont-PyMT mice: n=3, Trans-PyMT mice: n=4-7 from 3 independent experiments. **(b)** Similar activation of TBK1 in Spont- and Trans-PyMT tumors after DMXAA stimulation *in vitro*. Tumor cell suspensions from Spont- and Trans-PyMT mice were prepared as in Fig. 5b, cultured for 3 hours In vitro with DMXAA (250 µg/ml) or left untreated, then the levels of TBK1 activation (pTBK1) was determined by flow cytometry. The fold change in the percentage of $pTBK1^+$ cells in DMXA- versus control-treated cells was shown. Spont-PyMT mice: n=3, Trans-PyMT mice: n=8 . Data are from 3 independent experiments.

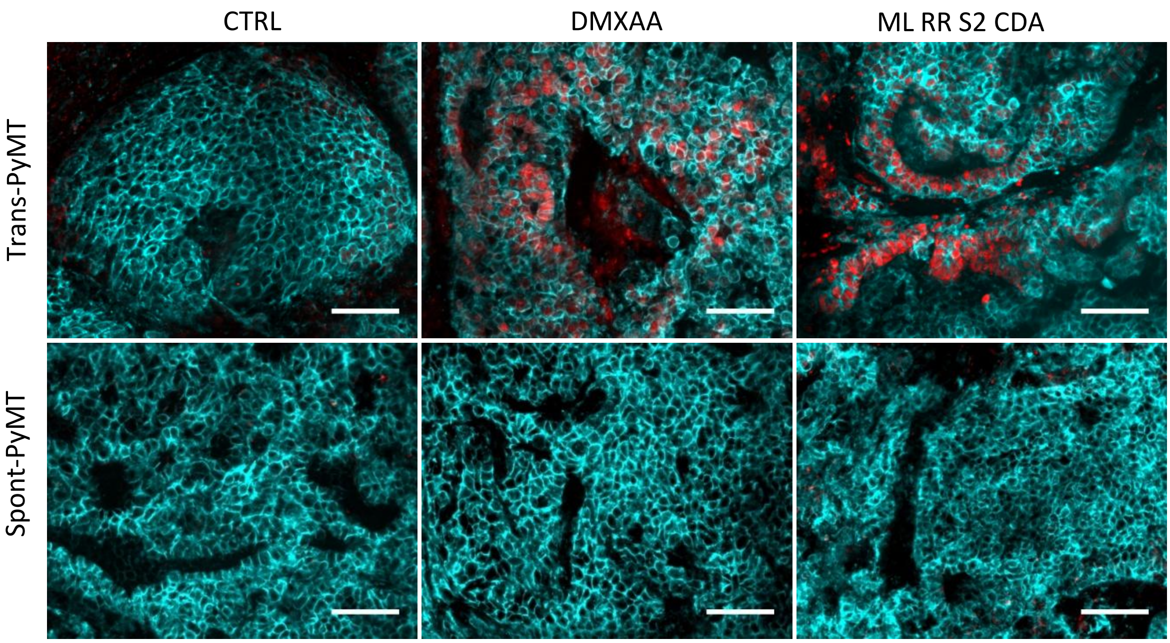


FigS3

**Figure S3. Intratumoral injection of DMXAA or ML RR-S2 CDA induced pIRF3 in Trans-PyMT mice, but not in Spont-PyMT mice. (a)** Trans- and Spont-PyMT mice were injected i.t. with DMXAA (1x 500µg) or ML RR-S2 CDA (1x 50µg). Tumors were collected 3 hours later and tumor slices stained with anti-EpCAM (blue) and anti-pIRF3 (red) mAbs. Scale bars = 50 µm. Images representative of Spont-PyMT mice: n=3 (CTRL), n=1 (DMXAA), n=2 (ML RR-S2 CDA); Trans-PyMT mice: n=2 (CTRL), n=2 (DMXAA), n=2 (ML RR-S2 CDA), from 5 independent experiments, are shown.

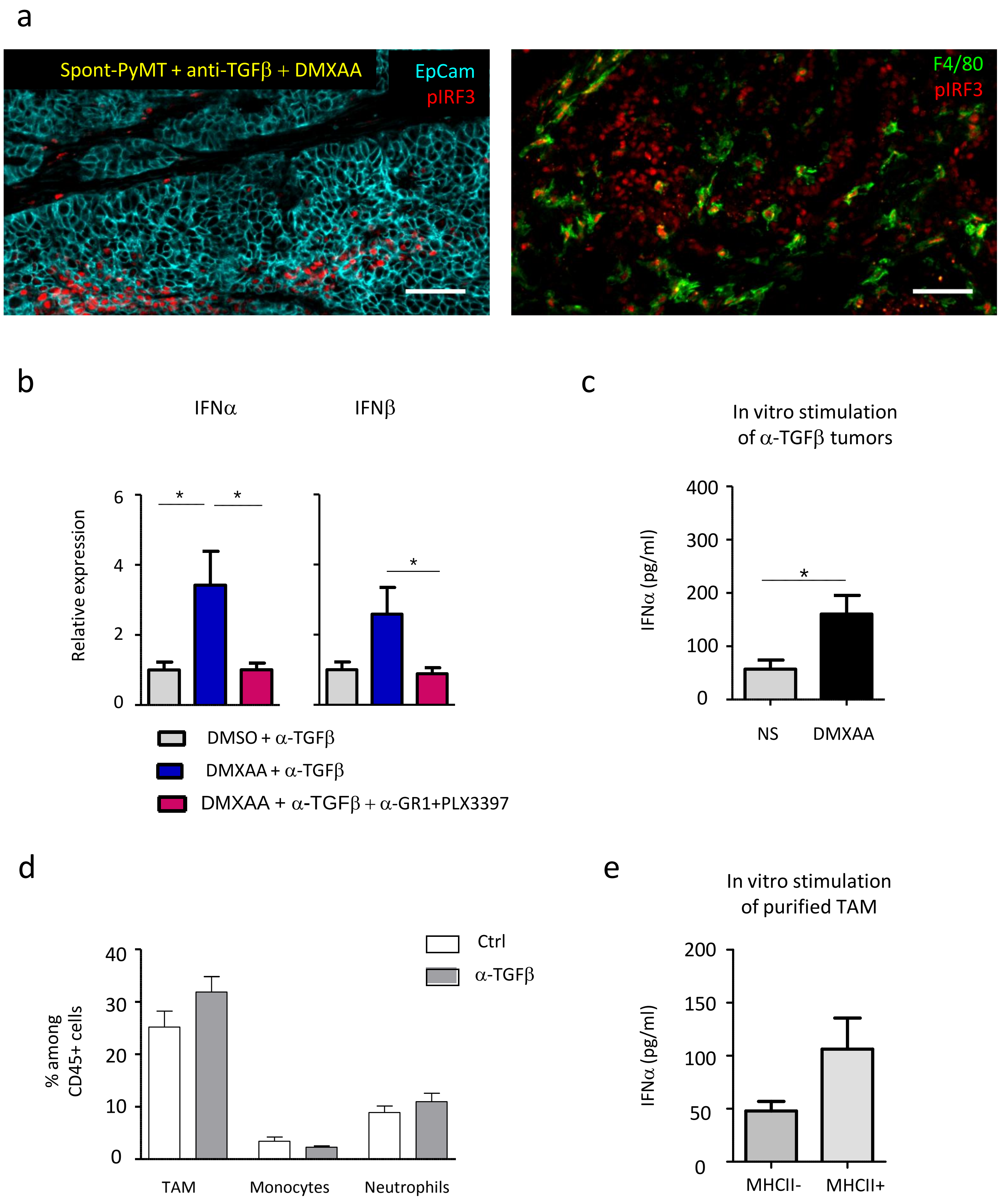

a
Spont-PyMT + anti-TGFβ + DMXAA
EpCam
pIRF3
F4/80
pIRF3
b
IFNα
IFNβ
Relative expression
0
2
4
6
*
*
*
DMSO + α-TGFβ
DMXAA + α-TGFβ
DMXAA + α-TGFβ + α-GR1+PLX3397
c
In vitro stimulation
of α-TGFβ tumors
IFNα (pg/ml)
0
100
200
300
400
*
NS
DMXAA
d
% among
CD45+ cells
0
10
20
30
40
Ctrl
α-TGFβ
TAM
Monocytes
Neutrophils
e
In vitro stimulation
of purified TAM
IFNα (pg/ml)
0
50
100
150
200
MHCII-
MHCII+


Fig S4

**Figure S4. Blocking TGFβ allows DMXAA-induced pIRF3 and type I IFN production in macrophages. (a)** Spont-PyMT mice, treated with an anti-TGFβ (day-4 and day-1), received one i.p. injection of DMXAA. Three hours later, tumors were taken and stained as in Fig. 5a. Two different image fields show that pIRF3+ cells may be tumor cells (EpCAM$^{+}$, blue, top), or myeloid cells (F4/80$^{+}$, green, bottom). Scale bars 50 µm. Images representative of *n*=4 treated Spont-PyMT mice. **(b)** Spont-PyMT mice, treated with an anti-TGFβ and anti-GR1 (day-4 and day-1) plus PLX3397 (from day-2 to day +1) received one i.p. injection of DMXAA (day0). One day later RNA from tumors were extracted to measure *Ifn*$\alpha$ and *Ifn*$\beta$ gene expression. The relative expression is shown, normalized to control tumors (DMSO, anti-TGFβ treated Spont-PyMT mice). Spont-PyMT mice treated with : DMSO+ anti-TGFβ (n=3), DMXAA+ anti-TGFβ (n=3); DMXAA+ anti-TGFβ+ anti-GR1 and PLX3397 (n=4) from 4 independent experiments. Results are expressed as mean ± s.e.m. **(c)** Tumors from anti-TGFβ treated Spont-PyMT mice were collected and pooled for each mouse, dissociated and restimulated *in vitro* with DMXAA (250µg/ml). IFNα produced in the supernatant after overnight culture was measured by ELISA. Results are expressed as mean ± s.e.m. Student *t*-Test. **(d)** Spont-PyMT mice were treated or not with an anti-TGFβ (day-4 and day-1) and sacrified at day 0. Tumors were dissociated and the proportion of TAM, Monocytes and neutrophils among CD45$^{+}$ cells was determined by flow cytometry. Spont-PyMT mice: *n*= 3, anti-TGFβ Spont-PyMT mice: *n*= 3. Data are from 3 independent experiments. Results are expressed as mean ± s.e.m. **(e)** MHC II$^{+}$ and MHC II$^{-}$ TAM from Spont-PyMT tumors were sorted by flow cytometry, stimulated overnight with DMXAA, and IFNα production was quantified by ELISA. Results are expressed as mean ± s.e.m. *n*=3 mice from 3 independent experiments.

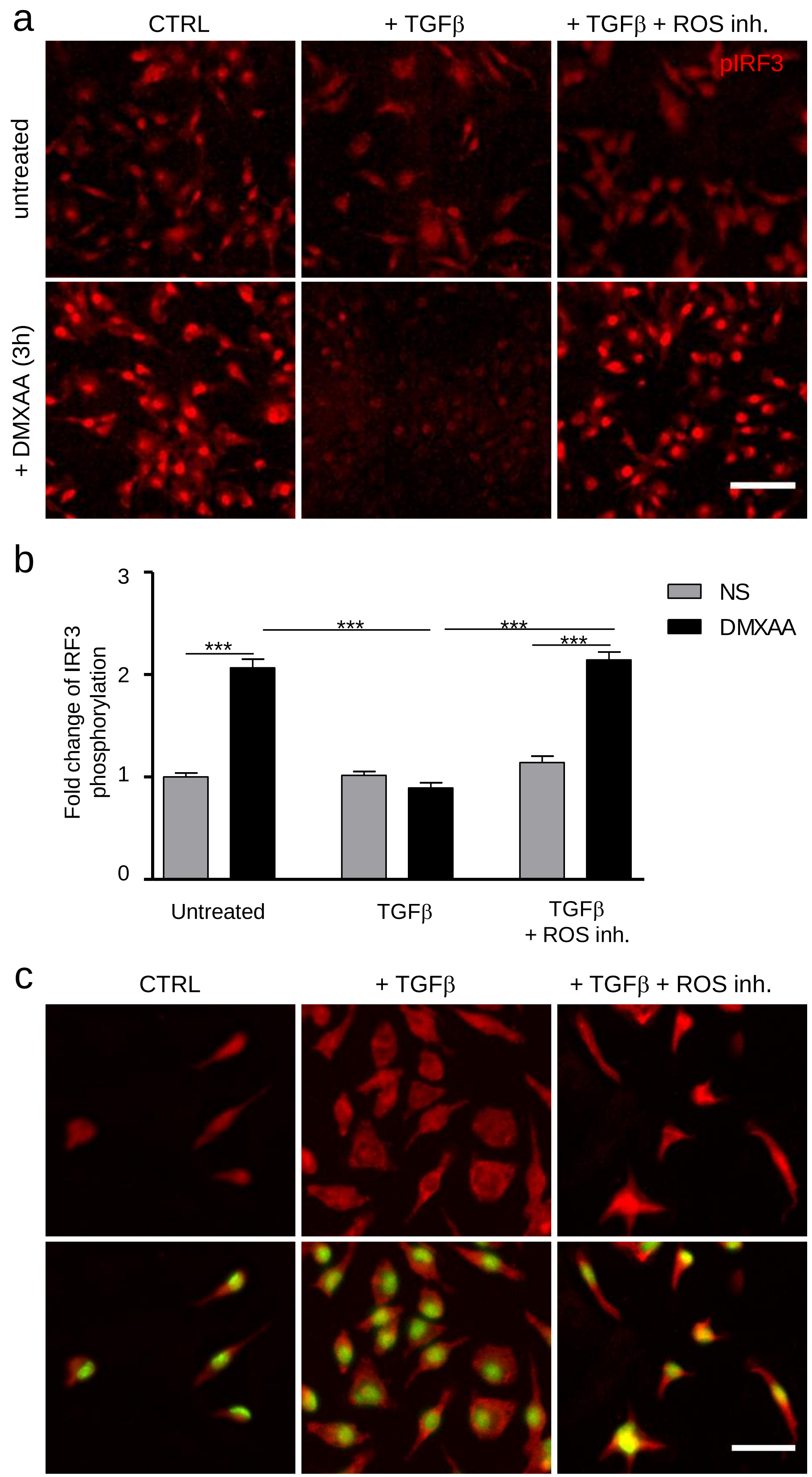
a
CTRL
+ TGFβ
+ TGFβ + ROS inh.
pIRF3
untreated
+ DMXAA (3h)
b
Fold change of IRF3 phosphorylation
3
2
1
0
***
***
***
***
NS
DMXAA
Untreated
TGFβ
TGFβ + ROS inh.
c
CTRL
+ TGFβ
+ TGFβ + ROS inh.

FigS5

**Figure S5. Exposure to TGFβ alters the capacity of BMDM to phosphorylate IRF3 after DMXAA stimulation and is associated with a ROS-dependent export of HDAC4 out of the nucleus. (a)** BMDM were cultured overnight with TGFβ (5ng/ml), TGFβ + ROS inhibitor (1nM) or left untreated. Cells were then stimulated or not with DMXAA (250µg/ml) for 3 hours, then stained with anti-pIRF3 Ab (red). Scale bar = 50 µm. **(b)** The increase in the surface covered by $pIRF3^+$ staining was measured in DMXAA-stimulated compared to untreated cells. Results are expressed as mean ± s.e.m from the cumulative data of 3 independent experiments. Tukey's Multiple Comparison Test. **(c)** BMDM exposed to TGFβ or TGFβ + ROS inhibitor as in **a**, were stained with anti-HDAC4 Ab (red) and, for the nucleus, DAPI (green). Top: HDAC4; Bottom: overlay with DAPI. Scale bar = 30 µm. Images from one experiment are shown and are representative of 3 independent experiments.

**Supplementary Table 1: Primer sequences for qPCR**

| | | |
|---|---|---|
| **IFNα4** | TGATGAGCTACTACTGGTCAGC | GATCTCTTAGCACAAGGATGGC |
| **IFNβ** | TGGGTGGAATGAGACTATTGTTG | CTCCCACGTCAATCTTTCTCT |
| **CCL2** | CATCCACGTGTTGGCTCA | GATCATCTTGCTGGTGAATGAGT |
| **CCL20** | GTGGGTTTCACAAGACAGATG | TTTTCACCCAGTTCTGCTTTG |
| **CXCL1** | GCTGGGATTCACCTCAATG | TGGGGACACCTTTTAGCATC |
| **TNFα** | AATGGCCTCCCTCTCATCAGTT | CGAATTTTGAGAAGATGATCTGAGTGT |
| **TGFβ** | TGACGTCACTGGAGTTGTACGG | GGTTCATGTCATGGATGGTGC |
| **GAPDH** | TGTGTCCGTCGTGGATCTGA | TTGCTGTTGAAGTCGCAG |

**Supplementary Table 2:** p values related to *in vivo* treatments shown in Fig. 6d after running a Kolmogorov-Smirnov test on the distribution of tumor growths across mice.

| | DMSO | DMSO + αTGFβ | DMXAA | DMXAA + αTGFβ | DMXAA + αTGFβ + αIFNaR | DMXAA + αTGFβ + αCD8 |
|---|---|---|---|---|---|---|
| DMSO | 1 | | | | | |
| DMSO + αTGFβ | 0.8668013 | 1 | | | | |
| DMXAA | 0.0790010 | 0.1608655 | 1 | | | |
| DMXAA + αTGFβ | 0.000003 | 0.0003630 | 0.00009 | 1 | | |
| DMXAA + αTGFβ + αIFNaR | 0.4167757 | 0.3180283 | 0.0486888 | 0.0003630 | 1 | |
| DMXAA + αTGFβ + αCD8 | 0.0001876 | 0.0023256 | 0.0018427 | 0.7794144 | 0.0023256 | 1 |
